# Harnessing Large Language Models for Structured Extraction of CYP–Substance Interactions from Biomedical Texts

**DOI:** 10.1101/2025.06.24.661414

**Authors:** Mariam Alkarmouty, Junya Ooka, Fumiyoshi Yamashita

## Abstract

Building on our previous work in biomedical text mining, we revisit the extraction of cytochrome P450 (CYP) and substance interactions using recent advances in large language models (LLMs). We present a scalable, high-accuracy framework that leverages the ChatGPT O3-mini model, employing optimized prompting and batch processing without relying on dictionaries or domain-specific ontologies. Our system achieves strong performance, with recall and precision of 0.950 and 0.978 across all CYP targets, and 0.923 and 0.993 for CYP3A4 specifically. This represents a substantial improvement over our earlier rule-based method. The resulting large-scale analysis not only reflects existing knowledge but also enables a more systematic and comprehensive integration of CYP isoform–substance interaction data, addressing the limitations of previous fragmented efforts. While previous studies have attempted to catalog these interactions, the scale, precision, and automation demonstrated here represent a significant step forward. These findings underscore the potential of LLM-driven pipelines to accelerate biomedical text mining and to support research in drug metabolism and related fields.

## 1. Introduction

Text mining has become a vital tool in biological and pharmaceutical research, enabling efficient extraction of knowledge from the growing body of scientific literature, patents, and clinical data (Krallinger et al., 2008). In drug discovery, it accelerates the identification of disease-related genes and drug targets, reduces costs, and supports drug repositioning and safety assessments (Tari and Patel, 2014).

In our previous work, we developed a natural language processing (NLP)-based text mining system to extract chemical–enzyme interactions from biomedical texts (Feng et al., 2007). Combining context-based and dictionary-based methods, the system achieved a recall of 0.852 and precision of 0.920, demonstrating high performance. Despite a remaining error rate of about 10%, it proved practical, allowing us to identify compounds interacting with cytochrome P450 (CYP) isoforms (Yamashita et al., 2011), pregnane X receptor (Yoshida et al., 2012), and P-glycoprotein (Yoshida et al., 2013), and to explore structure–activity relationships using extracted data.

However, difficulties such as linguistic variation and unfamiliar terminology revealed the limitations of existing methods and highlighted the need for more adaptive, semantically aware approaches. Large language models (LLMs), including ChatGPT, have recently gained attention for their ability to understand complex language through large-scale training (Li et al., 2024; Minaee et al., 2024), showing promise in tasks like scientific information retrieval and knowledge extraction (Gopalakrishnan et al., 2024; Hu et al., 2024; Bednarczyk et al., 2025).

While LLMs offer a potential alternative to rule-based systems (Gopalakrishnan et al., 2024; Hu et al., 2024; Bednarczyk et al., 2025), their performance in specialized domains like biomedical text mining—particularly for extracting chemical–biomolecule interactions—remains underexplored. Unlike traditional methods, LLMs can theoretically interpret diverse sentence structures and infer new concepts from context. However, prior studies suggest that for domain-specific tasks such as biomedical classification and causal relation extraction, general-purpose LLMs often underperform compared to fine-tuned models like BioBERT (Chen et al., 2024, 2025). Nevertheless, the landscape of LLMs is rapidly evolving, and it remains an open question whether recent models—such as GPT-4o and O3-mini—can overcome these limitations when applied to such specialized biomedical tasks.

In this study, we investigate the applicability of LLMs for extracting interaction data related to drug metabolism enzymes. By comparing the LLM-based method with our earlier system,^3^ we aim to assess its practical utility, scalability, and potential to support biomedical research and drug discovery.

## 2. Methods

### 2.1. Information Extraction From PubMed Abstracts

PubMed was searched for literature describing interactions between CYP enzymes and various substances. The abstracts of the retrieved articles were analyzed using the ChatGPT Batch API. The overall workflow is illustrated in Figure 1, and each step is described below.

**Figure 1.**
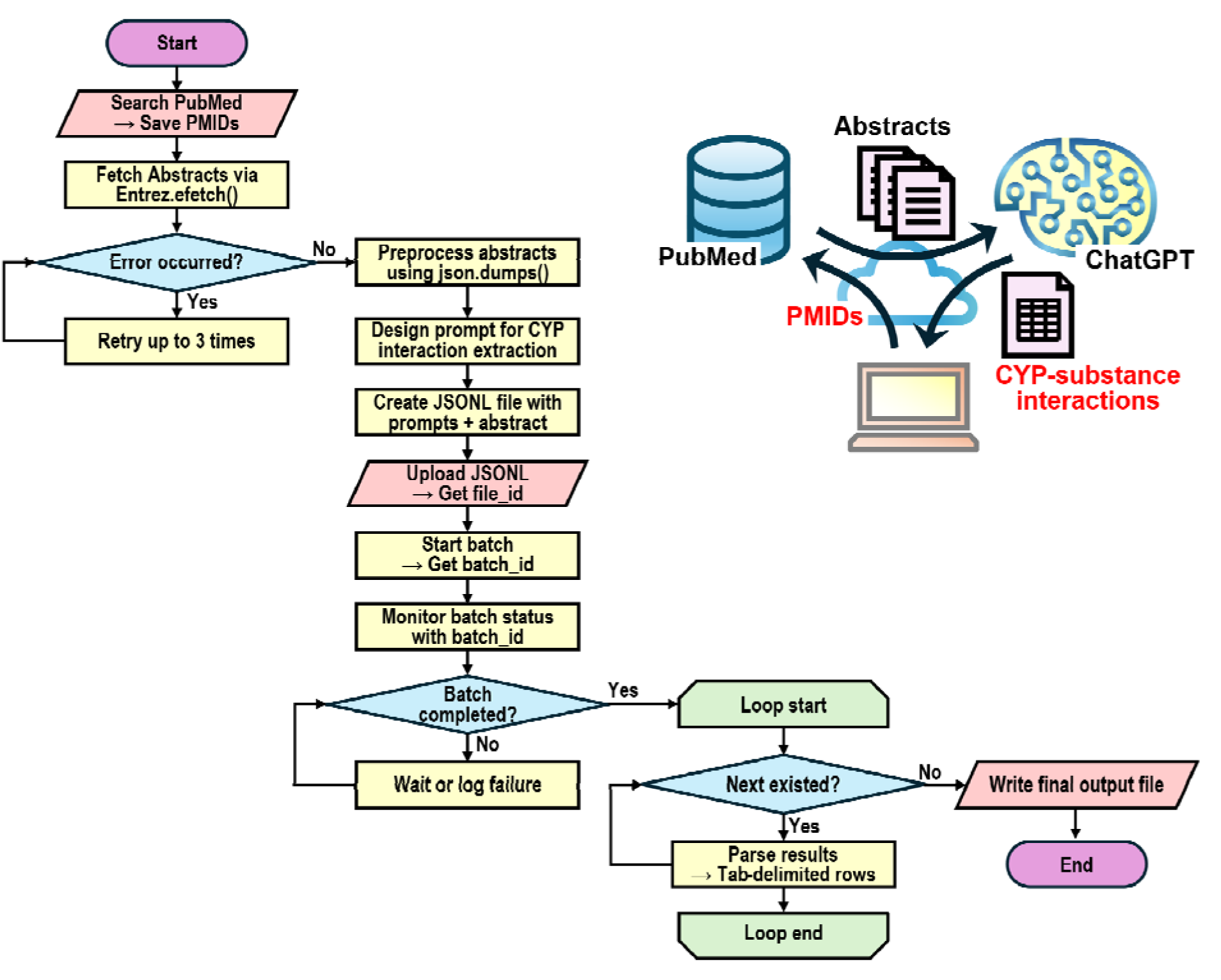
Workflow and schematic overview of ChatGPT Batch API–driven extraction of cytochrome P450–substance interaction information from PubMed abstracts.

#### 2.1.1. Preparation of input files for ChatGPT batch processing

A keyword search was conducted in PubMed, and the resulting PMIDs were saved to a text file. Using Biopython’s Entrez API, abstracts corresponding to each PMID were retrieved in XML format and parsed to extract relevant text.

The abstracts were converted into JSON-formatted strings compatible with the ChatGPT API. Prompts were designed to extract interaction types—substrate, inhibitor, inducer, or other—and the outputs were structured as tab-delimited tables (Table 1). Each API request included standardized instructions and definitions. The maximum token limit per completion was set to 4,000, and processing was carried out in batches of 1,000 entries.

**Table 1.** Prompt Structure for CYP–Substance Interaction Extraction Using ChatGPT Batch API.

| Role | Message |
| --- | --- |
| system | You are a helpful assistant. |
| user | From the given PubMed abstract, extract all interactions between cytochrome P450 isoforms and substances.<br>Instructions:<br>- Output a tab-delimited table without column headers. Use a **single tab character** (`'\t'`) to separate fields.<br>- **Do not** wrap the output in code blocks (e.g., triple backticks `````) or any other formatting.<br>- Output only the raw data rows — no explanations, no headings, no labels.<br>- Each row must represent one unique interaction between a cytochrome P450 isoform and a substance.<br>- For each interaction, include the following 4 fields, **in this exact order**:<br>1. Cytochrome P450 name<br>2. Substance name<br>3. Interaction type — select one of: substrate, inhibitor, inducer, or other<br>4. Mechanism of action (if specified; otherwise leave blank)<br>- If the abstract contains no interactions, return only: none<br>- Expand substance abbreviations only if their full form is defined in the abstract.<br>Definitions:<br>- Substrate: A substance metabolized by a cytochrome P450 enzyme.<br>- Inhibitor: A substance that reduces the activity of a cytochrome P450 enzyme.<br>- Inducer: A substance that increases the activity of a cytochrome P450 enzyme." |
| assistant | Please provide me the abstract. |
| user | (a PubMed abstract inserted by the program) |

All steps were implemented in Python v3.11, using libraries such as Biopython, requests, json, os, and time. The environment used was Google Colaboratory, with data stored on Google Drive. This setup enabled systematic processing of PubMed abstracts and the generation of structured data for downstream analysis (Listing S1).

#### 2.1.2. Batch submission and execution via the ChatGPT API

The prepared input JSONL files were submitted to the ChatGPT Batch API for processing (Listing S2). Each file was uploaded via a secure POST request to the OpenAI API. Upon receiving a file ID, batch processing was initiated by specifying the appropriate endpoint and a 24-hour execution window. To ensure reliability, a retry mechanism was applied to all API interactions. In the event of network issues or service errors, requests were retried up to three times. Batch IDs returned from successful submissions were recorded for tracking and reference.

#### 2.1.3. Batch monitoring and output processing

The final step involved monitoring batch status and processing the output (Listing S3). Batch statuses were checked through API calls. If a batch was marked “completed,” its output file ID was recorded for download.

Once no batches were in progress, completed output files were downloaded in JSONL format. Each line typically contained one abstract’s response with structured data. These responses were parsed to extract key fields, including the PubMed ID (embedded in a custom identifier) and the generated content. Each interaction was then converted into a tab-delimited format and written to a unified output file.

Common issues like missing fields or decoding errors were handled smoothly, ensuring the process could finish without interruption—even on large datasets. All API requests had built-in retry logic, automatically trying up to three times if temporary errors occurred, such as network issues or service unavailability.

### 2.2. Normalization and Standardization of Extracted Substance Names

Before normalization and metadata retrieval, a preprocessing step classified extracted chemical substance names. Using Excel’s *unique()* function, distinct names were collected for each CYP isoform and counted. This curated list served as input for the normalization pipeline (Listing S4). The workflow began by loading this list from a text file. Each name was standardized by cleaning special characters, unifying Greek letters, and removing extraneous symbols.

Normalized names were then matched against external databases to retrieve preferred names and structure-based identifiers through a three-stage process:

ChEMBL: Exact name matching to obtain molecular structures and standardized nomenclature.

PubChem: Queried if ChEMBL returned no result, using name- and structure-based lookups; InChIKeys were derived from desalinated SMILES.

UniProt: Used to identify full names for entries possibly referring to proteins or genes.

Multithreading was employed for efficiency. Matches were grouped by InChIKey, while unmatched entries were flagged for review. The final output—a tab-delimited file sorted by frequency—placed compounds lacking structural identifiers at the end for easier manual inspection.

### 2.3. Comparison with DrugBank Database

To enable direct matching with the text-mined interactions derived from PubMed abstracts, enzyme interaction data were first extracted from DrugBank 5.1.13 (Knox et al., 2024). An academic license was generously granted by the DrugBank team, providing free access to the full XML database. From this database, enzyme-related information was parsed for all listed drugs, with a specific focus on CYP enzymes. For each CYP enzyme associated with a given compound, the enzyme name and type of interaction (e.g., substrate, inhibitor, inducer) were recorded. To ensure consistency in classification, actions labeled as “downregulator” and “activator” in DrugBank were mapped to “inhibitor” and “inducer,” respectively.

To match chemical substances between DrugBank and the text-mined dataset, InChIKeys were used as unique identifiers, enabling precise structure-based normalization. Stereochemistry (chirality) was disregarded to account for naming variation and structural ambiguity. While this approach excluded biologics and certain natural products that lack associated InChIKeys, this limitation was considered acceptable, as CYP-mediated metabolism predominantly involves small-molecule compounds. Matching was performed separately for each CYP isoform and type of action.

## 3. Results and Discussion

### 3.1. Prompt Engineering for Structured Extraction of CYP-Substance Interactions

Prompt optimization was crucial for enabling structured and accurate extraction of cytochrome P450 (CYP)–substance interactions from biomedical literature using ChatGPT. Through iterative refinement, we developed a prompt that consistently generated tab-delimited outputs aligned with the target data schema (Table 1).

The prompt specified the use of a single tab character (\t) as the delimiter, with instructions to omit headers and code block formatting. These constraints ensured clean, machine-readable outputs suitable for downstream parsing.

To improve semantic consistency, brief definitions of interaction types—substrate, inhibitor, inducer, and other—were included, enabling the model to interpret varied terminology uniformly. Explicit instructions on output structure and ordering further enhanced reproducibility.

Importantly, the prompt directed the model to output only raw data rows, excluding explanatory text. Combined with preprocessing (e.g., escaping special characters using *json.dumps()*), this setup minimized formatting errors and ensured robust API communication during batch processing.

These design choices significantly improved data clarity and consistency, laying a solid foundation for subsequent validation and pharmacological analysis.

### 3.2. Large-Scale Extraction from PubMed Abstracts on CYP-Mediated Drug Metabolism

Using the keyword “cytochrome P450 drug metabolism” and filtering for “Abstract” (Text Availability), “English” (Language), and “Humans” (Species), we retrieved 33,986 abstracts from PubMed. Batch files of 1,000 requests each were generated based on their PMIDs and processed using the proposed system.

Of the 33,986 requests, 1,181 exceeded the maximum completion token limit (4,000 tokens). As no clear correlation between input and output token counts was observed (Figure S1), this was attributed to abstract complexity rather than length. Notably, the O3-mini model devoted an average of 96.4% of its completion tokens to reasoning content during processing, reflecting its reasoning-focused generation behavior (OpenAI, 2025). Given the relatively small number of over-limit cases and the cost of reanalysis, no retries were performed.

### 3.3. Benchmarking Text-Mining Models for CYP-Substance Interaction Extraction

To assess the performance of our system, we used a benchmark set of 100 CYP3A4-related abstracts that had previously been annotated for evaluating our earlier rule-based NLP system. Table 2 summarizes the recall and precision scores of various models on this dataset.

**Table 2.** Performance of information extraction by ChatGPT models^1)^

| Target | Model | Recall | Precision |
| --- | --- | --- | --- |
| All CYPs | O3-mini | 0.950 | 0.978 |
|  | GPT-4o | 0.774 | 0.872 |
|  | GPT-4o-mini | 0.771 | 0.785 |
| CYP3A4 | O3-mini | 0.923 | 0.993 |
|  | GPT-4o | 0.788 | 0.864 |
|  | GPT-4o-mini | 0.750 | 0.773 |
|  | our previous model<sup>2)</sup> | 0.852 | 0.920 |

- 1) The recall and precision of the models were evaluated using a set of 100 selected abstracts. Details validation results are presented in Supporting Information S2. - 2) The recall and precision of the NLP model we had previously developed were obtained from Feng et al. (2007).

O3-mini achieved the highest accuracy, with recall and precision of 0.950 and 0.978 across all CYPs, and 0.923 and 0.993 for CYP3A4 specifically. This represents a substantial improvement over our previous rule-based model, which achieved 0.852 and 0.920, respectively. In contrast, GPT-4o reached 0.774/0.872 and 0.788/0.864, while the lightweight GPT-4o-mini showed the lowest performance. These outcomes highlight the advantage of task-specific optimization embodied in O3-mini, which was designed to emphasize structured reasoning rather than long-context understanding or general efficiency (OpenAI, 2025). Despite GPT-4o’s larger capacity and superior context handling ability (OpenAI et al., 2024), it did not outperform O3-mini in this constrained information extraction task.

Recent benchmarks have shown that even state-of-the-art general-purpose models such as GPT-4 tend to underperform on biomedical information extraction tasks compared to fine-tuned domain-specific models. For instance, in the NCBI Disease dataset, BioBERT achieved an F1-score of 0.9090, whereas GPT-4 under zero-shot conditions reached only 0.5988—a difference of over 30 percentage points (Chen et al., 2025). While domain-specific models like BioBERT clearly have advantages, they require large-scale annotated data and substantial computational resources for fine-tuning, which fall outside the scope of this study.

Our objective was to explore the feasibility of general-purpose LLMs under zero- or few-shot conditions using only publicly accessible APIs. In this context, O3-mini demonstrated strong reliability and output consistency. Moreover, recent studies suggest that even high-performing models like GPT-4o exhibit weak calibration in biomedical settings, with minimal differences in confidence between correct and incorrect responses (Chen et al., 2024). While such limitations may be problematic in clinical decision-making, they are less critical in structured information extraction, where reproducibility and precision are more relevant.

Taken together, these findings support the use of prompt-optimized, lightweight LLMs for scalable biomedical text mining—particularly in use cases where simplicity, reproducibility, and low-cost deployment are prioritized over domain-specific fine-tuning.

### 3.4. Curation and Survey of CYP-Substance Interactions from Text Mining

A total of 65,165 CYP–substance interaction records were extracted via large-scale literature mining (Supporting Information S3). After removing duplicates, we retained only unique combinations of CYP isoform, substance name, and interaction type, yielding a curated dataset (Supporting Information S4). An overview of interaction patterns in this dataset is shown in Figure 2.

**Figure 2.**
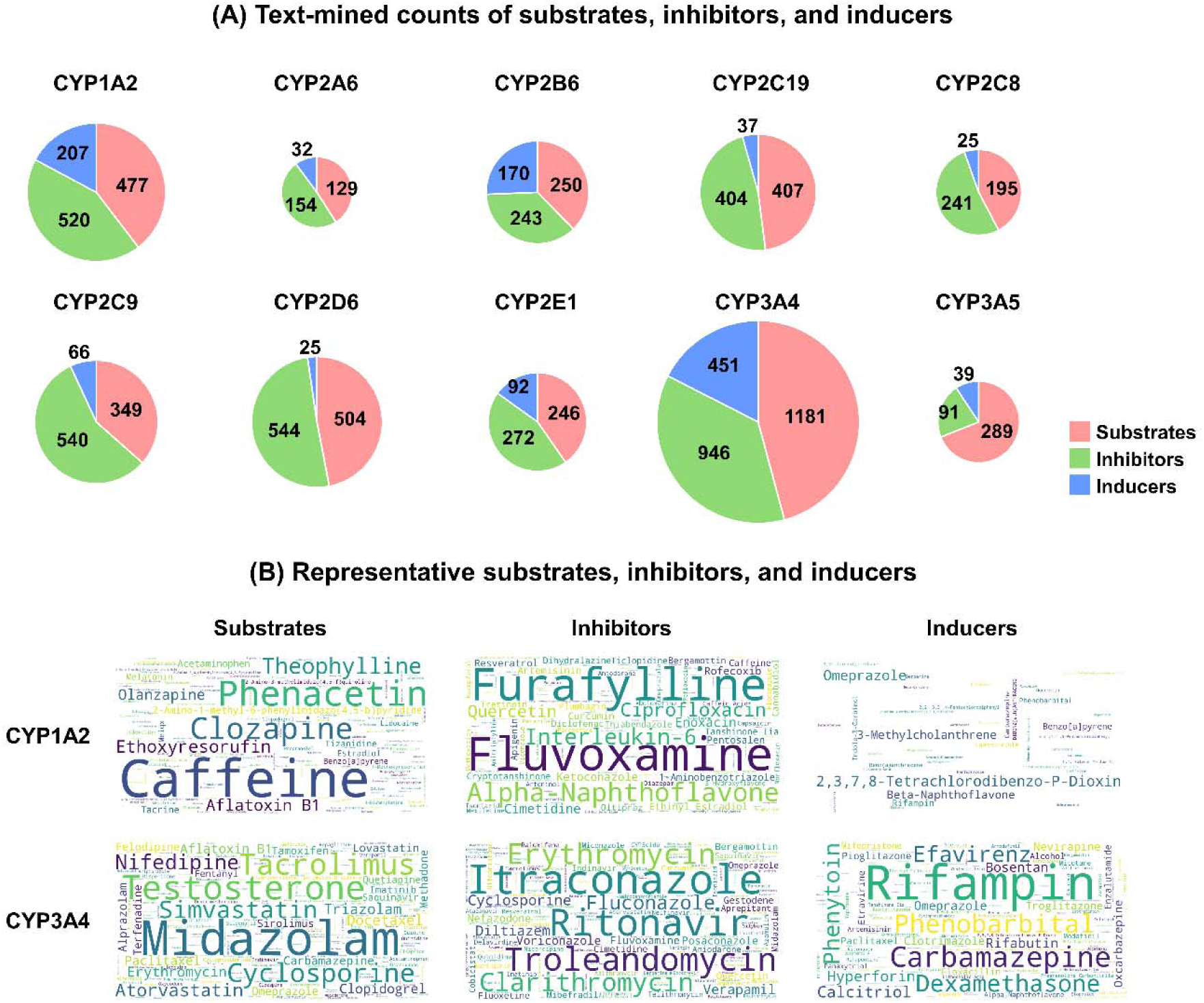
Summary and visualization of CYP-related substrates, inhibitors, and inducers identified via text mining. (A) Pie charts show the number of unique substances identified as substrates (green), inhibitors (red), or inducers (blue) for each major human cytochrome P450 (CYP) enzyme. (B) Representative substrates for each category are visualized as word clouds for CYP1A2 and CYP3A4. Word size reflects the frequency of occurrence in PubMed abstracts. Substrates, inhibitors, and inducers are shown in the left, center, and right panels, respectively.

Figure 2A illustrates key trends in the interaction profiles of major CYP isoforms. As expected, CYP3A4 emerged as the most prominent enzyme, with 2,578 recorded interactions— comprising 1,181 substrates, 946 inhibitors, and 451 inducers. This is consistent with its broad substrate specificity and critical role in drug–drug interactions (Zanger and Schwab, 2013). Other major isoforms, including CYP1A2, CYP2C9, CYP2C19, and CYP2D6, also exhibited substantial interaction counts. Notably, interaction patterns varied across isoforms: CYP2D6 interactions were predominantly substrates and inhibitors with few inducers, while CYP1A2 and CYP2E1 showed relatively higher inducer counts, reflecting their inducibility by environmental compounds.

Figure 2B presents a Wordcloud of representative compounds extracted by text mining. Frequently cited substances include established experimental probes such as caffeine and phenacetin for CYP1A2, and midazolam and testosterone for CYP3A4. Additionally, well-known strong inhibitors like fluvoxamine and itraconazole, along with inducers such as rifampin and omeprazole, were prominently featured.

### 3.5. Comparison with DrugBank-Derived Interaction Data

To assess the pharmacological relevance and completeness of the text-mined interactions, we compared our curated dataset with enzyme–compound associations listed in DrugBank (Knox et al., 2024). Matching was performed based on InChIKeys, ignoring stereochemistry, as it is not consistently reported in PubMed abstracts. Entries without InChIKeys, such as biologics and certain natural products, were excluded. This is a reasonable limitation, given that CYP metabolism predominantly involves small molecules.

Figure 3 (substrates) and Supplementary Figure S2 (inhibitors and inducers) illustrate the overlap between CYP-associated compounds identified in DrugBank and those obtained through text mining of PubMed abstracts. For nearly all CYP isoforms and interaction types, the number of compounds retrieved by our approach exceeded those listed in DrugBank. This reflects the broader scope of literature-based extraction, which is not limited to approved or investigational drugs. Among the 2,978 structurally defined compounds extracted in this study, 1,518 (approximately 51%) were also found in DrugBank, while the remainder consisted of non-drug entities such as dietary components, environmental contaminants, endogenous metabolites, and laboratory tool compounds.

**Figure 3.**
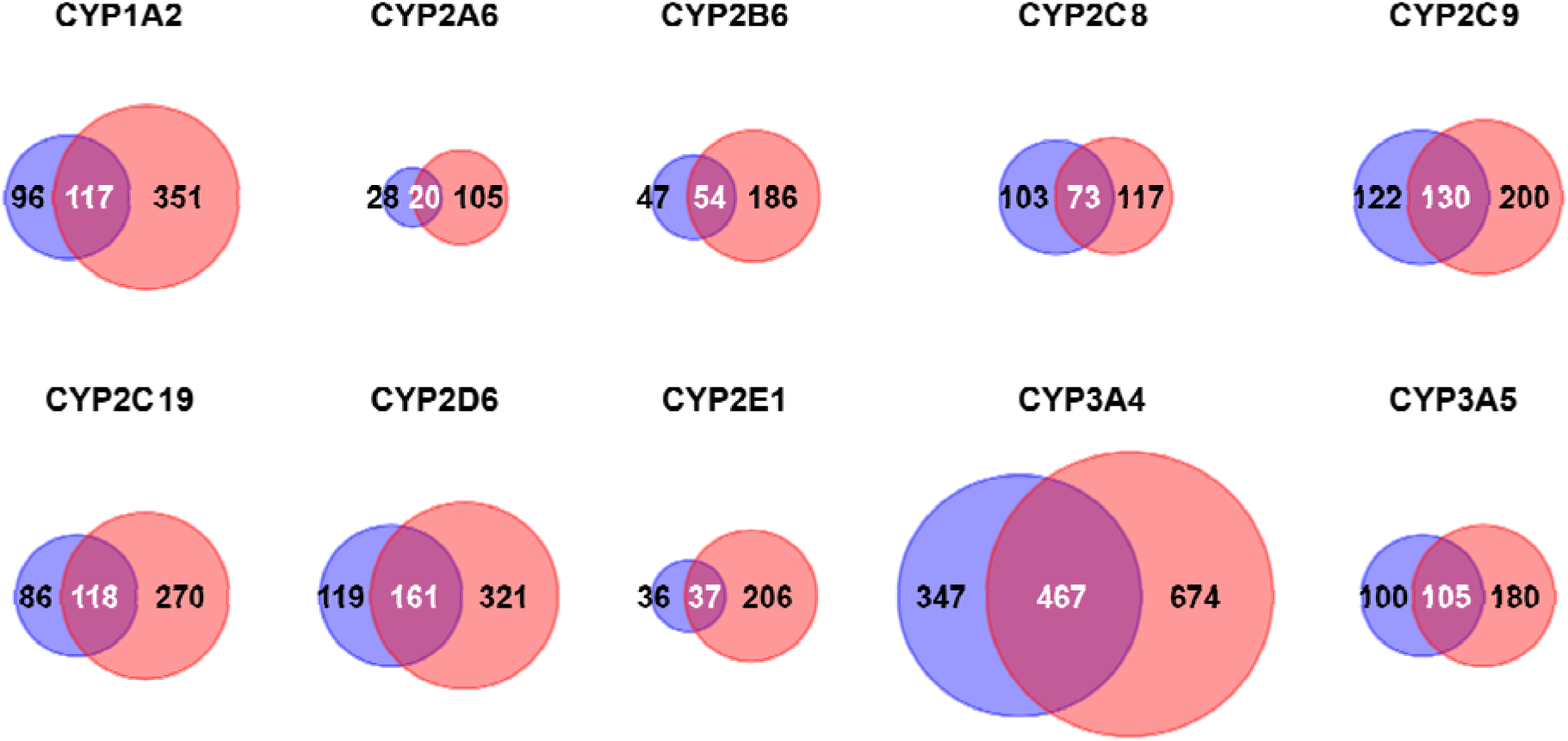
Comparison of Text-Mined CYP Substrates with Those Listed in DrugBank. Venn diagrams show the number of unique CYP substrates extracted from PubMed abstracts (red) and listed in DrugBank (blue). Substance matching was based on InChIKeys, ignoring stereochemistry. Only structurally defined small molecules were included.

In the curated dataset provided as Supplementary Material S4, each CYP interaction is color-coded to indicate whether the compound is listed in DrugBank and whether the interaction is consistent with DrugBank annotations. Agreement with DrugBank is fairly high for substrates. However, our text mining approach identified a substantial number of inhibitors and inducers that are not annotated in DrugBank, despite the compounds themselves being listed. This highlights potential gaps in the coverage of curated databases, particularly for interaction types that are more context-dependent or less well characterized. One possible reason for this discrepancy is that our approach tends to include positive findings even when evidence is conflicting. At the same time, the differences reflect distinct curation strategies. For example, the case of lansoprazole illustrates one of the strengths of DrugBank’s conservative annotation policy. Although lansoprazole has been reported as an inducer of CYP1A2, it is not annotated as such in DrugBank, likely because induction does not occur under conventional therapeutic conditions (Dilger et al., 1999; Rizzo et al., 1996). This cautious approach helps maintain the reliability of curated data by minimizing overinterpretation of uncertain or context-specific findings.

Nevertheless, our analysis also indicates that literature mining can uncover a variety of interactions that are not yet incorporated into DrugBank. For instance, in the metabolism of arachidonic acid, CYP2C8 has been shown to play a more prominent role than CYP2C9 (Rifkind et al., 1995; Dai et al., 2001), yet only CYP2C9 is listed. Similarly, although primaquine is primarily metabolized by CYP2D6 (Baird et al., 2018; Potter et al., 2015), it is annotated only as an inhibitor. Extensive literature also supports the involvement of CYP2A6 in the human metabolism of coumarin (Pelkonen et al., 2000), but the compound is described solely as a CYP2C9 substrate. In other cases, such as 2-amino-1-methyl-6-phenylimidazo(4,5-b)pyridine (PhIP) (Teunissen et al., 2010) and indole-3-carbinol (Reed et al., 2005), the absence of metabolic annotation may simply reflect areas where curation has not yet caught up with the literature.

Conversely, there are also compounds listed in DrugBank that were not captured by our text mining approach, representing false negatives. This shortfall can be attributed to several factors. First, DrugBank integrates information from a wide range of sources, including full-text articles, drug labels, and regulatory documents, while our method relies exclusively on PubMed abstracts. As a result, well-established interactions that are not explicitly mentioned in abstracts may be missed. Second, although our method achieved a high precision of approximately 99%, the recall was around 92%, indicating that some relevant interactions may not be detected even within the available abstracts.

Taken together, these observations suggest that integrating literature-mined data with existing curated databases may help provide a more comprehensive and up-to-date understanding of CYP-related drug interactions.

## 4. Conclusion

The proposed system leverages prompt-optimized LLMs to enable accurate and structured extraction of cytochrome P450–substance interactions from biomedical literature at scale. By combining well-engineered prompts with robust batch processing and validation workflows, the system achieved strong performance in both general extraction and isoform-specific analysis. Our large-scale results not only aligned with existing knowledge but also enabled a more systematic and comprehensive collection of CYP isoform–substance interaction data, helping to consolidate previously fragmented information. This large-scale extraction provides a valuable resource on CYP-mediated metabolism, supporting drug development, drug–drug interaction risk assessment, and the identification of candidate compounds for further in vitro or in silico studies. These findings demonstrate the reliability of our approach and highlight the potential of LLM-based pipelines to advance biomedical text mining, offering scalable synthesis of domain-specific knowledge without the need for manual curation. The methodology presented here has potential applications beyond drug metabolism, across a wide range of biomedical research domains.

## CRediT authorship contribution statement

Mariam Alkarmouty: investigation, resources, writing – review & editing. Junya Ooka: investigation, resources, writing – review & editing. Fumiyoshi Yamashita: conceptualization, investigation, methodology, resources, writing – original draft, writing – review & editing, funding acquisition, and project administration.

## Declaration of competing interest

The authors declare that they have no known competing financial interests.

## Disclosure of generative AI usage

The authors utilized ChatGPT-4o to check and refine the grammar, syntax, and clarity of the manuscript.

## Acknowledgements

This study was supported by grants from the Japan Society for the Promotion of Science (JSPS) KAKENHI (grant numbers: 23H02650 (FY) and 23K17475 (FY)). We would also like to thank the DrugBank Team for granting us an academic license, which greatly facilitated our research.

## Supporting Information

**Supporting Information S1.** Relationship between input tokens and completion tokens (Figure S1); Comparison of Text-Mined CYP Inhibitors and Inducers with DrugBank Entries (Figure S2); Script to Generate Input Data for ChatGPT Batch Processing (Listing S1); Script to Upload JSON-Formatted Batch Files and Start ChatGPT Batch Jobs (Listing S2); Script to Retrieve and Parse ChatGPT Batch Results (Listing S3); Script to Normalize and Annotate Chemical Names (Listing S4) (DOCX).

**Supporting Information S2.** Summary of the text-mining performance of various models (XLSX).

**Supporting Information S3.** Raw extracted data of CYP–substance interactions (XLSX). **Supporting Information S4.** Curated CYP–substance interaction data for representative CYP isoforms (XLSX).

## Data Availability

We provided the program scripts for text mining, as well as the raw data extracted and the processed data, as Supporting Information. All other data used in this study are available at: https://osf.io/zt29b/.

## Notes

### Competing Interest Statement

The authors have declared no competing interest.

