## Supporting Information S1 for "Harnessing Large Language Models for Structured Extraction of CYP–Substance Interactions from Biomedical Texts"

***Supplementary File 1***

**Harnessing Large Language Models for Structured Extraction of CYP–Substance Interactions from Biomedical Texts**

Mariam Alkarmouty and Fumiyoshi Yamashita*

Department of Quantitative Pharmaceutics, Graduate School of Pharmaceutical Sciences, Kyoto University, Sakyo-ku, Kyoto 606-8501, Japan.

**Contents**

**Figure S1.** Relationship between input tokens and completion tokens

**Figure S2.** Comparison of Text-Mined CYP Inhibitors and Inducers with DrugBank Entries

**Listing S1.** Script to Generate Input Data for ChatGPT Batch Processing

**Listing S2.** Script to Upload JSON-Formatted Batch Files and Start ChatGPT Batch Jobs

**Listing S3.** Script to Retrieve and Parse ChatGPT Batch Results

**Listing S4.** Script to Normalize and Annotate Chemical Names

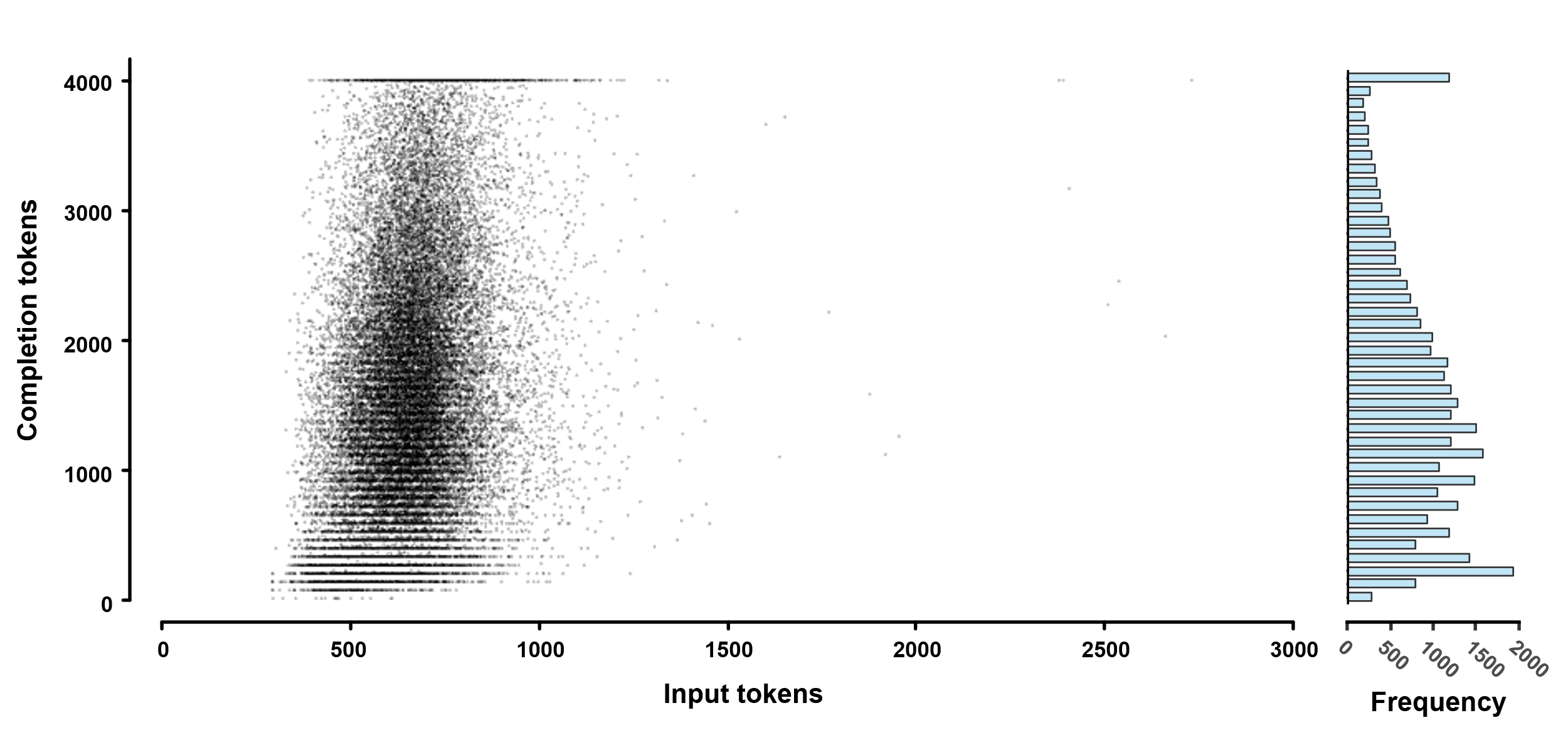

**Figure S1.** Relationship between input tokens and completion tokens.

No correlation between input tokens and completion tokens was observed. This aligns with O3-mini’s reasoning-first design: OpenAI notes that the model is “trained to think for longer before responding,” relying on a long internal chain of thought before it begins to write, so output length is determined by answer complexity rather than prompt size.^1,2^ Consequently, the O3-mini model consumed, on average, 96.4 % of its allocated completion tokens during reasoning.

Refs.

1) https://openai.com/index/introducing-o3-and-o4-mini/

2) https://platform.openai.com/docs/guides/reasoning?utm_source=chatgpt.com&api-mode=chat

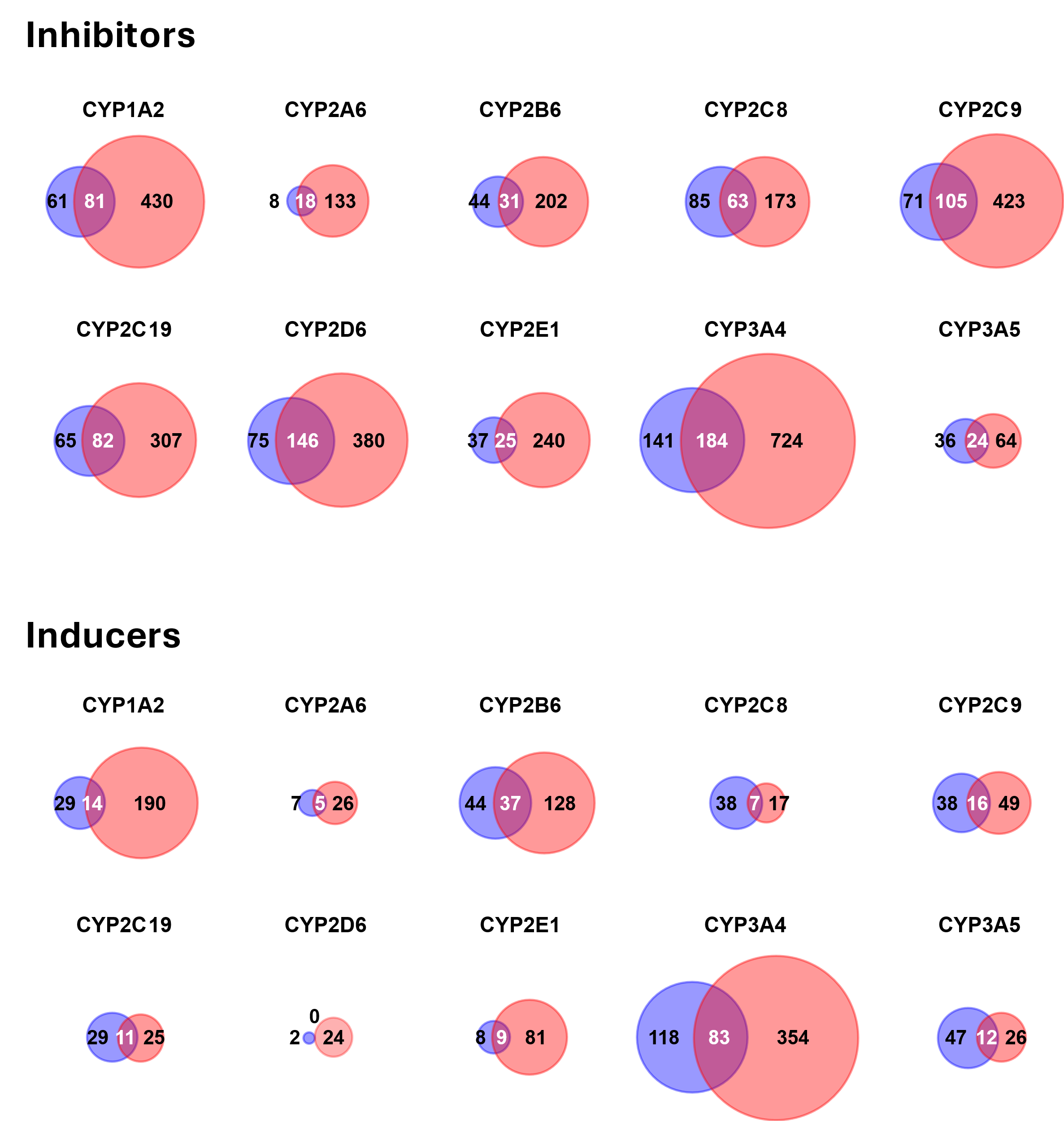

**Figure S2.** Comparison of Text-Mined CYP Inhibitors and Inducers with DrugBank Entries.

Venn diagrams show the number of unique CYP substrates extracted from PubMed abstracts (red) and listed in DrugBank (blue). Substance matching was based on InChIKeys, ignoring stereochemistry. Only structurally defined small molecules were included.

**Listing S1.** Script to Generate Input Data for ChatGPT Batch Processing

```
!pip install biopython

from Bio import Entrez
from time import sleep
import os
from google.colab import drive

# --- Configuration ---
Entrez.email = ""
INTERVAL_YEARS = 5
START_YEAR = 1960
END_YEAR = 2024
MAX_RECORDS = 10000
RETMAX = 10000
SLEEP_TIME = 0.34

QUERY_BASE = '"human cytochrome p450 drug metabolism" AND hasabstract[text] AND english[lang]'
OUTPUT_DIR = "/content/drive/My Drive/CypTextAnalysis"
OUTPUT_FILE = "pmids.txt"
OUTPUT_PATH = os.path.join(OUTPUT_DIR, OUTPUT_FILE)

# Mount Google Drive
drive.mount('/content/drive')
os.makedirs(OUTPUT_DIR, exist_ok=True)

def get_year_ranges(start, end, interval):
    return [(y, min(y + interval - 1, end)) for y in range(start, end + 1, interval)]

def fetch_pmids_for_range(start_y, end_y):
    query = f'{QUERY_BASE} AND ("{start_y}"[PDAT] : "{end_y}"[PDAT])'
    handle = Entrez.esearch(db="pubmed", term=query, usehistory="y", retmax=0)
    result = Entrez.read(handle)
    handle.close()

    count = int(result["Count"])
    print(f"  → Found {count} PMIDs")
    if count > MAX_RECORDS:
        raise RuntimeError(f"Too many results in {start_y}–{end_y} ({count}). Reduce interval.")

    pmids = []
    webenv = result["WebEnv"]
    query_key = result["QueryKey"]

    for start in range(0, count, RETMAX):
        print(f"    Fetching {start + 1}–{min(start + RETMAX, count)}")
        handle = Entrez.esearch(
            db="pubmed",
            webenv=webenv,
            query_key=query_key,
            retstart=start,
            retmax=RETMAX,
            retmode="xml"
        )
        data = Entrez.read(handle)
        handle.close()
        pmids.extend(data["IdList"])
        sleep(SLEEP_TIME)
    return pmids

# --- Main Process ---
all_pmids = []
for start_y, end_y in get_year_ranges(START_YEAR, END_YEAR, INTERVAL_YEARS):
    print(f"\nSearching {start_y}–{end_y}")
    all_pmids.extend(fetch_pmids_for_range(start_y, end_y))

# Deduplicate and save
unique_pmids = list(dict.fromkeys(all_pmids))
with open(OUTPUT_PATH, "w") as f:
    f.write("\n".join(unique_pmids))

print(f"\nDone! {len(unique_pmids)} unique PMIDs saved to: {OUTPUT_PATH}")
```

**Listing S2.** Script to Upload JSON-Formatted Batch Files and Start ChatGPT Batch Jobs

```
import os
import time
import json
import requests
from google.colab import drive

# --- Configuration ---
API_KEY = "sk-proj-***"  # OpenAI API Key
BASE_URL = "https://api.openai.com/v1"
HEADERS = {'Authorization': f'Bearer {API_KEY}'}
DRIVE_PATH = "/content/drive"
FOLDER_PATH = os.path.join(DRIVE_PATH, "My Drive/CypTextAnalysis")
INPUT_FILE = "chunk_json_list.txt"
OUTPUT_FILE = "batch_id_list.txt"

# --- Initialize session ---
session = requests.Session()
session.headers.update(HEADERS)

# --- Utility functions ---
def request_with_retry(method, url, **kwargs):
    """Send HTTP request with retry logic."""
    for attempt in range(3):
        try:
            response = session.request(method, url, **kwargs)
            if response.ok:
                return response.json()
            print(f"[HTTP {response.status_code}] {response.text}")
        except Exception as e:
            print(f"Request error (attempt {attempt + 1}): {e}")
        time.sleep(2)
    return None

def upload_file(filepath, purpose="batch"):
    """Upload a file to OpenAI for batch processing."""
    if not os.path.exists(filepath):
        print(f"File not found: {filepath}")
        return None

    with open(filepath, 'rb') as f:
        files = {'file': f, 'purpose': (None, purpose)}
        return request_with_retry("POST", f"{BASE_URL}/files", files=files)

def start_batch(file_id):
    """Start batch processing with the given file ID."""
    data = {
        'input_file_id': file_id,
        'endpoint': '/v1/chat/completions',
        'completion_window': '24h'
    }
    res = request_with_retry("POST", f"{BASE_URL}/batches", json=data)
    return res.get("id") if res else None

def process_files():
    """Upload input files and start batch jobs. Save batch IDs."""
    input_path = os.path.join(FOLDER_PATH, INPUT_FILE)
    output_path = os.path.join(FOLDER_PATH, OUTPUT_FILE)

    if not os.path.exists(input_path):
        print(f"Input file not found: {input_path}")
        return

    with open(input_path, "r", encoding="utf-8") as f:
        filenames = [line.strip() for line in f if line.strip()]

    if not filenames:
        print("No filenames found in input.")
        return

    with open(output_path, "a", encoding="utf-8") as out_f:
        for name in filenames:
            filepath = os.path.join(FOLDER_PATH, name)
            file_res = upload_file(filepath)

            if file_res and (file_id := file_res.get("id")):
                batch_id = start_batch(file_id)
                if batch_id:
                    print(f"Uploaded: {name}, Batch ID: {batch_id}")
                    out_f.write(batch_id + "\n")
                else:
                    print(f"Failed to start batch for: {name}")
            else:
                print(f"Failed to upload file: {name}")
            time.sleep(180)  # Optional: delay for rate limit compliance

# --- Execute ---
if __name__ == "__main__":
    drive.mount(DRIVE_PATH, force_remount=True)
    os.makedirs(FOLDER_PATH, exist_ok=True)

    process_files()
```

**Listing S3.** Script to Retrieve and Parse ChatGPT Batch Results

```
import os
import json
import time
import requests
from google.colab import drive

# --- Configuration ---
API_KEY = "sk-proj-***"  # OpenAI API Key
BASE_URL = "https://api.openai.com/v1"
HEADERS = {'Authorization': f'Bearer {API_KEY}'}
DRIVE_PATH = "/content/drive"
FOLDER_PATH = os.path.join(DRIVE_PATH, "My Drive/CypTextAnalysis")
INPUT_FILE = "batch_id_list.txt"
OUTPUT_FILE = "result.txt"

# --- Initialize session ---
session = requests.Session()
session.headers.update(HEADERS)

# --- Utility functions ---
def request_with_retry(method, url, **kwargs):
    """Send HTTP request with retry logic."""
    for attempt in range(3):
        try:
            response = session.request(method, url, **kwargs)
            if response.ok:
                return response.json()
            print(f"[HTTP {response.status_code}] {response.text}")
        except Exception as e:
            print(f"Request error (attempt {attempt + 1}): {e}")
        time.sleep(2)
    return None

def check_batch_status(batch_id):
    """Return (status, output_file_id) for a given batch ID."""
    res = request_with_retry("GET", f"{BASE_URL}/batches/{batch_id}")
    if not res:
        print(f"Failed to retrieve status for batch: {batch_id}")
        return None, None
    return res.get("status"), res.get("output_file_id")

def download_results(file_id):
    """Download output file content from OpenAI."""
    try:
        response = session.get(f"{BASE_URL}/files/{file_id}/content")
        if response.ok:
            return response.text
        print(f"Download failed for file {file_id} (status {response.status_code})")
    except Exception as e:
        print(f"Download error for file {file_id}: {e}")
    return None

def parse_and_write_results(response_text, output_file):
    """Parse OpenAI batch response and write formatted result."""
    for line in response_text.splitlines():
        time.sleep(0.02)  # Throttle for readability or rate-limiting
        try:
            json_obj = json.loads(line)
            pmid = json_obj.get("custom_id", "unknown").split("-")[1] if "custom_id" in json_obj else "unknown"
            choices = json_obj.get("response", {}).get("body", {}).get("choices", [])
            if choices:
                content = choices[0].get("message", {}).get("content", "")
                for entry in content.splitlines():
                    clean_entry = entry.replace('\\t', '\t')
                    output_file.write(f"{pmid}\t{clean_entry}\n")
        except (json.JSONDecodeError, IndexError) as e:
            print(f"Error parsing line: {e}")

# --- Main process ---
def main():
    drive.mount(DRIVE_PATH, force_remount=True)
    os.makedirs(FOLDER_PATH, exist_ok=True)

    input_path = os.path.join(FOLDER_PATH, INPUT_FILE)
    output_path = os.path.join(FOLDER_PATH, OUTPUT_FILE)

    if not os.path.exists(input_path):
        print(f"Input file not found: {input_path}")
        return

    with open(input_path, "r", encoding="utf-8") as f:
        batch_ids = [line.strip() for line in f if line.strip()]

    output_ids = []
    for batch_id in batch_ids:
        status, file_id = check_batch_status(batch_id)
        if status == "completed":
            output_ids.append(file_id)
        elif status == "in_progress":
            print(f"Batch {batch_id} is still in progress.")
            return
        else:
            print(f"Batch {batch_id} has failed or expired.")

    if not output_ids:
        print("No completed batch outputs found.")
        return

    with open(output_path, "w", encoding="utf-8") as out_file:
        for file_id in output_ids:
            result_text = download_results(file_id)
            if result_text:
                parse_and_write_results(result_text, out_file)

    print(f"Results written to: {output_path}")

# --- Execute ---
if __name__ == "__main__":
    main()
```

**Listing S4.** Script to Normalize and Annotate Chemical Names

```
!pip install rdkit chembl-webresource-client -q

import os
import re
import time
import json
import unicodedata
import requests
import pandas as pd
from difflib import SequenceMatcher
from concurrent.futures import ThreadPoolExecutor
from urllib3.util.retry import Retry
from requests.adapters import HTTPAdapter

from rdkit import Chem
from rdkit.Chem.MolStandardize import rdMolStandardize
from chembl_webresource_client.new_client import new_client
from google.colab import drive

# --- Google Drive Setup ---
drive.mount('/content/drive', force_remount=True)
PATH_FOLDER = "/content/drive/My Drive/ColabNotebooks"
os.makedirs(PATH_FOLDER, exist_ok=True)

# --- HTTP Session Setup ---
session = requests.Session()
retries = Retry(total=5, backoff_factor=2, status_forcelist=[429, 500, 502, 503, 504])
adapter = HTTPAdapter(max_retries=retries)
session.mount("http://", adapter)
session.mount("https://", adapter)

# --- Normalize chemical/protein name ---
def normalize_name_string(name):
    name = unicodedata.normalize('NFKC', name).lower()
    greek_map = {
        "α": "alpha", "β": "beta", "γ": "gamma", "δ": "delta", "ε": "epsilon",
        "ζ": "zeta", "η": "eta", "θ": "theta", "ι": "iota", "κ": "kappa",
        "λ": "lambda", "μ": "mu", "ν": "nu", "ξ": "xi", "ο": "omicron",
        "π": "pi", "ρ": "rho", "σ": "sigma", "ς": "sigma", "τ": "tau",
        "υ": "upsilon", "φ": "phi", "χ": "chi", "ψ": "psi", "ω": "omega"
    }
    for sym, word in greek_map.items():
        name = name.replace(sym, word)

    name = re.sub(r"[’‘´`]", "'", name)
    name = re.sub(r"[®©™]", "", name)
    name = re.sub(r"[-–−]+", "-", name)
    name = re.sub(r"[^\w\s,\+\-\(\)\[\]']", "", name)
    name = re.sub(r"\s+", " ", name).strip()
    name = re.sub(r"^-+|--+", "-", name)
    name = re.sub(r'\s\([^)]+\)$', '', name)
    return name

# --- Score if a name is likely trivial ---
def is_likely_trivial_name(title):
    if not title:
        return 0
    score = 0
    if len(title) <= 15:
        score += 2
    if not re.search(r'[\d\-,()=]', title):
        score += 2
    if re.fullmatch(r'[A-Za-z ]+', title):
        score += 2
    if title[0].islower():
        score += 1
    return score

# --- Query PubChem for compound info ---
def get_pubchem_info(query, by="name"):
    time.sleep(0.25)
    try:
        if by == "iupac":
            res = session.get(f'https://opsin.ch.cam.ac.uk/opsin/{query}.json', timeout=10)
            res.raise_for_status()
            inchikey = res.json().get('stdinchikey')
            if not inchikey:
                return None
            query, by = inchikey, "inchikey"

        if by in {"name", "inchikey"}:
            res = session.get(f'https://pubchem.ncbi.nlm.nih.gov/rest/pug/compound/{by}/{query}/cids/JSON', timeout=10)
            if res.status_code == 404:
                return None
            res.raise_for_status()
            cids = list(set(res.json().get("IdentifierList", {}).get("CID", [])))
            if not cids:
                return None
            query, by = cids, "cid"

        if isinstance(query, list):
            best_match, best_score = None, -1
            for cid in query:
                res = session.get(f"https://pubchem.ncbi.nlm.nih.gov/rest/pug/compound/cid/{cid}/property/InChIKey,Title,SMILES/JSON", timeout=10)
                if res.status_code == 404:
                    continue
                res.raise_for_status()
                prop = res.json()["PropertyTable"]["Properties"][0]
                score = is_likely_trivial_name(prop.get("Title", ""))
                if score > best_score:
                    best_score = score
                    best_match = prop
            if best_match:
                return {
                    "inchikey": best_match.get("InChIKey", ""),
                    "pref_name": best_match.get("Title", ""),
                    "smiles": best_match.get("SMILES", "")
                }
        return None
    except Exception as e:
        print(f"[PubChem Error] {by}: {query} | {str(e)}")
        return None

# --- Query UniProt for protein info ---
def query_uniprot_name(input_name):
    base_url = "https://rest.uniprot.org/uniprotkb/search"
    exclude = ["receptor", "binding", "associated", "subunit", "like", "regulatory"]
    if any(term in input_name.lower() for term in exclude):
        return None
    exclude_clause = " ".join([f'AND NOT protein_name:{term}' for term in exclude])
    query = f'protein_name:"{input_name}" AND organism_id:9606 AND reviewed:true {exclude_clause}'
    params = {
        "query": query,
        "format": "json",
        "fields": "accession,id,protein_name,organism_name,length",
        "size": 1
    }
    try:
        res = requests.get(base_url, params=params, timeout=10)
        res.raise_for_status()
        results = res.json().get("results", [])
        if not results:
            return None
        result = results[0]
        return {
            "UniProt Accession": result["primaryAccession"],
            "Entry Name": result["uniProtkbId"],
            "Protein Name": result["proteinDescription"]["recommendedName"]["fullName"]["value"],
            "Organism": result["organism"]["scientificName"],
            "Length": result["sequence"]["length"]
        }
    except Exception as e:
        print(f"[UniProt Error] {input_name} | {str(e)}")
        return None

# --- Convert SMILES to desalted InChIKey ---
def desalt_to_inchikey(smiles):
    mol = Chem.MolFromSmiles(smiles)
    if not mol:
        return None
    parent = rdMolStandardize.LargestFragmentChooser().choose(mol)
    return Chem.MolToInchiKey(parent) if parent else None

# --- Normalize name and collect metadata ---
def normalize_name(name_count_tuple):
    name, count = name_count_tuple
    cleaned = normalize_name_string(name)

    def make_result(pref_name, inchikey, flag="success"):
        return {
            "input_name": name,
            "desalted_pref_name": pref_name,
            "desalted_inchikey": inchikey,
            "count": count,
            "flag": flag
        }

    if len(cleaned) <= 3:
        return make_result("Too short to analyze", "Too short", flag="failure")

    try:
        chembl_hits = list(new_client.molecule.filter(pref_name__iexact=cleaned))
        if chembl_hits:
            mol = chembl_hits[0]
            struct = mol.get("molecule_structures", {})
            pref_name = mol.get("pref_name", "").title()

            if struct:
                smiles = struct.get("canonical_smiles")
                inchikey = struct.get("standard_inchi_key")
                if smiles and inchikey:
                    desalted = desalt_to_inchikey(smiles) or inchikey
                    if desalted != inchikey:
                        info = get_pubchem_info(desalted, by="inchikey")
                        if info:
                            return make_result(info["pref_name"], info["inchikey"])
                    return make_result(pref_name, inchikey)

            if pref_name:
                return make_result(pref_name, "Not applicable")

        for by in ["name", "iupac"]:
            info = get_pubchem_info(cleaned, by=by)
            if info:
                smiles = info["smiles"]
                desalted = desalt_to_inchikey(smiles) or info["inchikey"]
                info = get_pubchem_info(desalted, by="inchikey") if desalted != smiles else info
                return make_result(info["pref_name"], info["inchikey"])

        protein = query_uniprot_name(cleaned)
        if protein:
            return make_result(protein["Protein Name"], "Not applicable")

    except Exception as e:
        print(f"[Normalization Error] {name} | {str(e)}")

    return make_result("Not found", "Not available", flag="failure")

# --- Load input and normalize ---
input_path = os.path.join(PATH_FOLDER, "chemicals_list.txt")
if os.path.exists(input_path):
    with open(input_path, 'r', encoding='utf-8') as f:
        name_count_pairs = [(name, int(count)) for name, count in (line.strip().split('\t') for line in f)]
else:
    print("File not found!")
    exit()

with ThreadPoolExecutor(max_workers=8) as executor:
    results = list(executor.map(normalize_name, name_count_pairs))

df = pd.DataFrame(results)

# --- Group and prepare output ---
df_valid = df[df['flag'] == 'success']
df_invalid = df[df['flag'] == 'failure'][['input_name', 'desalted_inchikey', 'desalted_pref_name', 'count', 'flag']]

idx = df_valid.groupby('desalted_inchikey')['count'].idxmax()
rows_with_max_count = df_valid.loc[idx]
sum_counts = df_valid.groupby('desalted_inchikey', as_index=False)['count'].sum()
df_grouped = rows_with_max_count.drop(columns='count').merge(sum_counts, on='desalted_inchikey')

df_final = pd.concat([df_grouped, df_invalid], ignore_index=True)
df_output = df_final[df_final['flag'] == 'success'].copy()
df_output['inchikey_na_flag'] = df_output['desalted_inchikey'] == 'Not applicable'

df_output_sorted = df_output.sort_values(
    by=['inchikey_na_flag', 'count'],
    ascending=[True, False]
).rename(columns={'count': 'sum_counts'})

output_path = os.path.join(PATH_FOLDER, "output_prefname_count.txt")
df_output_sorted[['input_name', 'desalted_inchikey', 'desalted_pref_name', 'sum_counts']].to_csv(
    output_path, sep='\t', index=False, encoding='shift_jis'
)

# --- Summary Output ---
print(f"\nResults saved to: {output_path}")
```
